# Evaluation of porcine epidemic diarrhea virus RNA contamination on swine industry transportation vehicles

**DOI:** 10.1101/2025.01.24.634714

**Authors:** Taylor B. Parker, Kelly A. Meiklejohn, Gustavo Machado, Michael Rahe, Bradford Sean Darrow, Juliana Bonin Ferreira

**Affiliations:** Department of Population Health and Pathobiology, North Carolina State University, 1060 William Moore Drive, Raleigh, NC, 27606 USA

**Keywords:** swine, PEDV, RNA detection, between farm virus transmission, RT-qPCR, vehicle cleaning and disinfection, glutaraldehyde, quaternary ammonium

## Abstract

Porcine epidemic diarrhea virus (PEDV) is one of the most devastating pathogens of global swine health and welfare. It is well known that contaminated fomites and vehicle movements play an important role in farm-to-farm PEDV spread, but the efficacy of cleaning and disinfection (C&D) protocols on the reduction in dissemination risk via vehicles and trailers remains unclear. This study used swine industry data to determine how frequently vehicles and trailers were contaminated with PEDV RNA before and after C&D. Environmental RNA samples were collected at three eastern North Carolina C&D sites from four different vehicle types: crew trucks, feed trucks, pigs-to-farm trucks and trailers, and pigs-to-market trucks and trailers. A total of 2,004 samples were collected from truck cabins, trailers, and tires before and after C&D with two commercial disinfectants at two different concentrations. An in-house RT-qPCR assay was used to detect the presence of PEDV RNA only (not infectivity status). Results suggest that pigs-to-market trucks hauling live pigs were the most likely to be contaminated with PEDV (82.06% of trucks tested positive before C&D and 89.55% tested positive after C&D), while feed trucks were the least likely contaminated (11.73% of trucks testing positive before C&D and 11.25% testing positive after C&D). Based on PEDV RNA detection, we demonstrated that quaternary ammonium and glutaraldehyde is a more effective disinfectant compared to advanced hydrogen peroxide in eliminating detectable PEDV RNA. Results also show that truck cabins are just as contaminated as the exterior of their vehicles. Based on these results, vehicle biosecurity measures should be evaluated and modified to prevent the spread of PEDV.

## 1. Introduction

Porcine epidemic diarrhea virus (PEDV) is one of the most devastating pathogens of global swine health and welfare. This highly contagious enteric pathogen of pigs causes vomiting, diarrhea, and dehydration in swine of all ages, as well as high mortality in neonatal piglets (Stevenson et al., 2013; Lei et al., 2024). PEDV was introduced into the U.S. swine industry in 2013 when 16 states reported a total of 218 cases within the first nine weeks (Stevenson et al., 2013; Mole, 2013; Lowe et al., 2014; Lee, 2015; Machado et al., 2019). It is now estimated that roughly 55 - 60% of U.S. commercial breeding herds have been exposed to the virus at one time or another (AASV, 2013; Goede et al., 2015; Kikuti et al., 2022). Moreover, evidence from field veterinarians shows that PEDV remains ranked among the top concerns from veterinary and production standpoints (VanderWall and Deen, 2018). Over a decade since its introduction into the U.S., PEDV continues to be an important issue for the industry due to the economic burden from both direct and indirect costs associated with the occurrence of an outbreak, which can reach up to USD $300,000 (based on the closure of a 700-head breeding herd) (Weng et al., 2016). As of April 2024, the Morrison Swine Health Monitoring Project (MSHMP) reported an incidence of 3.8% of PEDV cases in U.S. sow herds (MSHMP Chart 1). This has been an improvement since 2015, but the virus still costs the U.S. swine industry over USD $50 million per year (National Hog Farmer, 2024).

PEDV can be transmitted directly through the fecal-oral route, as well as via contaminated fomites (Stevenson et al., 2013; Bowman et al., 2015; Lei et al., 2024; Houston et al., 2024). It is also known that vehicle movements play a role in disease spread (Lee et al., 2019; Lowe et al., 2014; Boniotti et al., 2018). However, the importance of pathogen transmission via truck cabins, live-haul trailers, feed trucks, and crew trucks in the swine industry is under-researched (Stevenson et al., 2013; Houston et al., 2024). Biosecurity measures including decontamination via cleaning and disinfecting (C&D) protocols are in place to help reduce virus transmission, but studies have shown that these methods are not always effective (Boniotti et al., 2018; Houston et al., 2024; Li et al., 2020). Therefore, understanding how vehicle movement and decontamination affect disease outbreaks is crucial to decreasing the transmission of PEDV (Galvis and Machado, 2024).

The objective of this study was to use swine industry data to determine how frequently swine production vehicles were contaminated with PEDV RNA before and after C&D following the industry SOP which included washing (including soaking and removal of organic material) and disinfection. While addressing this question, the effectiveness of selected commercial disinfectants at eliminating detectable PEDV RNA, as well as how PEDV RNA contamination varied across truck classifications and sampling locations (e.g., interior of truck cabin vs. trailer or tire), were evaluated. Notably, this current study did not assess whether PEDV RNA detected on swine vehicles was capable of causing infection.

## 2. Materials and Methods

Figure 1 provides a schematic of the key steps in this study’s experimental design, which are described in detail in the following subsections.

**Figure 1.**
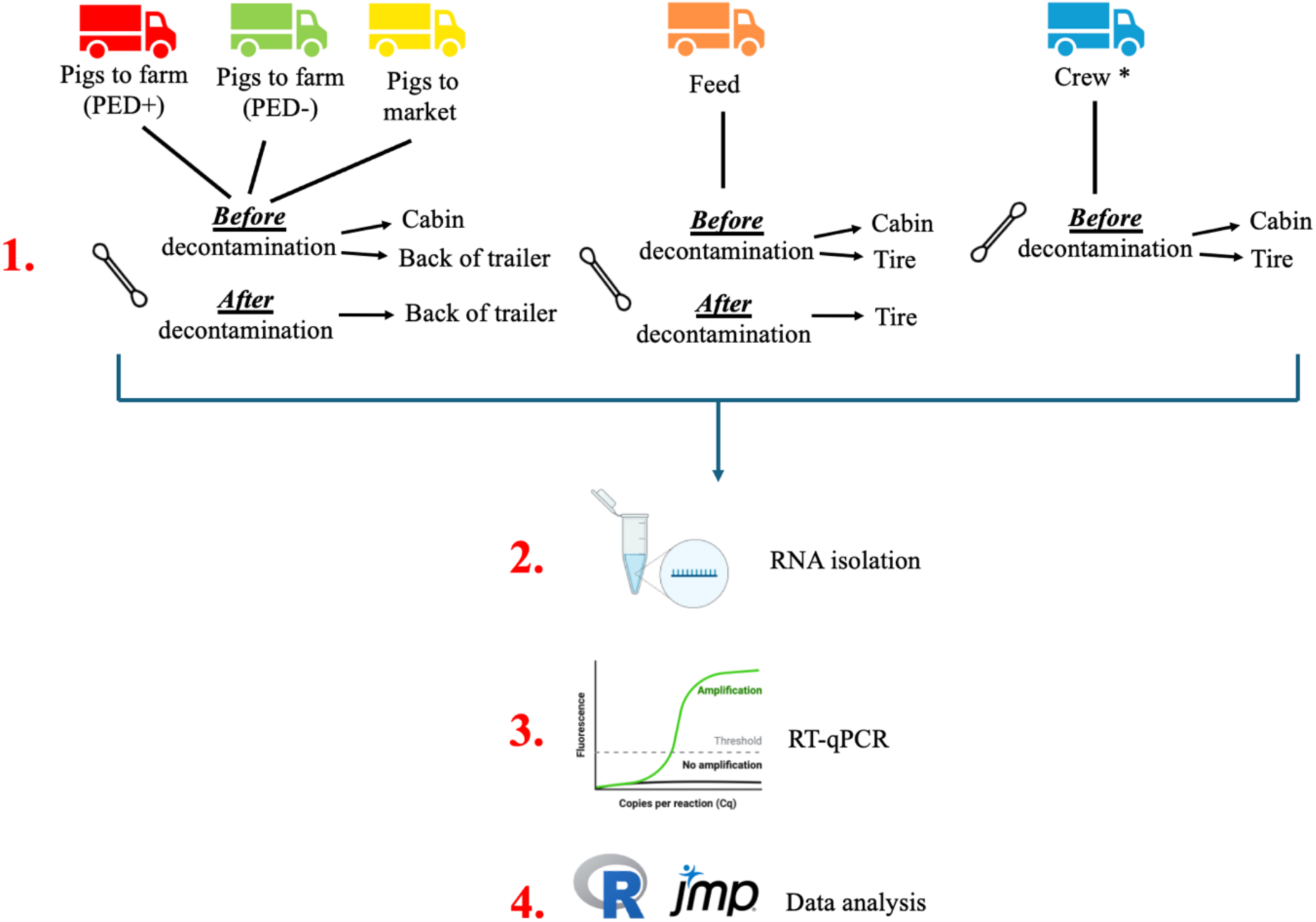
Sampling and processing workflow used in this study. * denotes that crew trucks were not decontaminated at C&D stations, therefore “after decontamination” samples were not collected.

### 2.1 Selection of truck types and disinfectants

A total of four different truck types used by the participating commercial swine company were sampled in this study: 1) pigs-to-farm trucks, which transport live pigs between farms (trucks are classified depending on whether they are coming from farms that were experiencing a PEDV outbreak [referred to as “PED+ trucks”] and trucks coming from farms that were not experiencing a PEDV outbreak [referred to as “PED-trucks”]), 2) pigs-to-market trucks, which transport pigs to markets (i.e. slaughterhouse, packing plants; referred to as “market trucks”), 3) feed trucks, which carry feed from the feed mill to farms, and 4) crew trucks, which are used to move personnel performing a wide range of farm tasks (e.g. vaccination, power washing at closeouts, pig loading, and unloading). Pigs-to-farm and market trailers were power washed by the swine company personnel to remove debris and organic material, then disinfectant was sprayed onto to the inside and outside of the trailers. The lower portion of feed trucks (e.g. tires and surfaces close to the ground) are sprayed with disinfectant in between loads and crew trucks are washed using a company car wash after daily shifts. The sample size for each truck type was determined using a power analysis with a 95% confidence interval assuming 30% of the selected 398 trucks were positive for PEDV (*pers comm*). Trucks were sampled at three specific C&D stations located in eastern North Carolina, U.S. selected due to: a) locality close to a high hog density area, and b) surrounding herds have a history of ongoing PEDV. All C&D stations used a unidirectional flow pattern and recycled water for washing. For confidentiality, stations will be numbered/identified as 1, 2, and 3 (Table 1).

**Table 1:** Description of truck washes included in this study and corresponding truck types.

| C&D station | Number of bays for C&D | Active disinfectant ingredient | Disinfectant dilutions studied | Truck types (month[s] of sampling) |
| --- | --- | --- | --- | --- |
| 1 | 1* | Accelerated Hydrogen Peroxide | 1:32 and 1:40 | Pigs-to-market (Nov-Feb); Pigs-to-farm (PED-) (Mar-May) |
| 2 | 1 | Quaternary Ammonium and Glutaraldehyde | 1:100 and 1:128 | Pigs-to-farm (PED+) (Nov-May) |
| 3 | 3 | Quaternary Ammonium and Glutaraldehyde | 1:100 and 1:128 | Pigs-to-farm (PED-) (Nov-Feb); Pigs-to-market (Mar-May); feed. |

\* denotes there are three other bays at this station that utilized Quaternary Ammonium and Glutaraldehyde, but were not included in this study.

The rationale for disinfectant choice was based on those currently used by the swine industry and field veterinarians, and what was available at the three selected C&D stations. The production company running the C&D stations typically disinfects using quaternary ammonium and glutaraldehyde (QAG) at a 1:256 dilution and advanced hydrogen peroxide (AHP) at a 1:32 dilution. However, previous studies have compared different agricultural disinfectants and found discordant results regarding the efficacy of their ability to lower the concentration of PEDV RNA (Bowman et al., 2015; Holtkamp et al., 2016; Houston et al., 2024). Given this, we opted to test both disinfectants at two concentrations each. All three C&D stations rotated their disinfectant concentration between a higher and lower concentration for each collection period (Table 1). For QAG, the higher concentration was diluted at 1:100 and the lower concentration was diluted at 1:128. For AHP, the higher concentration was diluted at 1:32 and the lower was diluted at 1:40. While these were the stated concentrations, it was not possible to internally confirm a) if the working disinfectant solutions were prepared to these concentrations by truck wash workers (however, C&D station personnel verified the disinfectant concentration after preparing according to their validated procedure), and b) the volume of disinfectant administered to each truck. To measure the working concentrations of both disinfectants, test strips (QAG– Cid Lines, Ypres, Belgium; AHP– Virox Technologies Inc., Oakville, ON, Canada) were used to test the concentration of disinfectant after application to the back of the trailers of PED-pigs-to-farm and pigs-to market trucks.

The disinfectants in use at each of the C&D stations were as follows (Table 1): *Station 1* - QAG used in three bays (not sampled in this study) and AHP in the fourth bay per our request; *Station 2* - QAG in single bay; *Station 3* - QAG in all three bays, one of which was dedicated to feed truck disinfection. In March, the production company opted to move PED-pigs-to-farm trucks to C&D station 1 and pigs-to-market trucks to C&D station 3. Crew trucks were washed at the end of shifts at a drive-through car wash operated by the production company and therefore not subjected to the included disinfectants.

### 2.2 Sample Collection

For each of the four truck types, both an interior cabin sample and exterior truck/trailer sample were collected per vehicle. The exterior sample was collected at either the trailer or the tire (for vehicle types that did not carry live animals). Specific collection sites for each truck type were as follows: 1) *pigs-to-farm and market trucks*, the cabin floorboard underneath the brake pedal and the loading zone of the trailer and surrounding hinges; 2) *feed trucks*, the cabin floorboard underneath the brake pedal and one tire from the feed-holding trailer; and 3) *crew trucks*, the cabin floorboard underneath the brake pedal and one tire from the truck (Figure 1). When collecting a sample, a Puritan Hydraflock swab (Puritan Medical Products, Guilford, ME, USA) was moistened by dipping into a 1.5 mL collection tube containing 500 µL DNA/RNA Shield (Zymo Research Corporation, Irvine, CA, USA) and then subsequently wiped and rolled across approximately a 20-by-20 cm area before placing the swab head back into the collection tube. The first truck for each truck type from each bi-weekly collection period was swabbed in triplicate; two samples served as collection duplicates and a third sample was used as a “processing positive control” (described below). Samples were kept on ice during collection and stored at -80°C upon arrival at the laboratory. Samples were collected bi-weekly from 27 Nov 2023 to 31 May 2024 to include peak PEDV season which has a higher incidence from late fall to spring.

Additionally, to act as a positive control, samples were collected via swab from various locations at a farm confirmed to be PEDV+ (e.g. pen wall and floor, boot sole, and feed trough) and stored in 500 µL of DNA/RNA Shield. Upon arrival at the laboratory, samples were frozen at -80°C.

### 2.3 Detection of PEDV

Reverse transcription-quantitative PCR (RT-qPCR) is a useful molecular biology method that allows researchers to detect specific sequences of RNA in a sample using custom nucleic acid primers that bind to and amplify a target sequence. Once amplified to a detectable level, a fluorescent dye can be measured which correlates to the quantity of the amplified RNA of the target sequence in the sample. Commercial RT-qPCR assays are available for the detection of many swine viruses, but are often expensive and not tailored to a specific sample type. This study uses a custom in-house RT-qPCR assay that was designed to detect PEDV RNA specifically from environmental vehicle samples. Sections 2.3.1 and 2.3.2 detail the specific methods and reagents used in this assay.

#### 2.3.1 RNA Isolation

The positive control samples collected from the PED+ farm were thawed at room temperature and isolated using the Quick-DNA/RNA Viral Kit (Zymo Research Corporation) according to the manufacturer’s instructions. Eluates across multiple RNA isolations were pooled, separated into 500 µL aliquots, and stored at -80°C until use.

The environmental samples collected from trucks were thawed to room temperature and swabs were placed in spin baskets set inside the collection tube (ChromeTech of Wisconsin Inc, Franklin, WI, USA) and centrifuged at 12,000 rpm for two minutes. This step was completed to a) ensure complete recovery of all liquid retained in the swab head, and b) facilitate the removal of large debris and particulates adhered to the swab. A total of 375 µL of supernatant was transferred into a 1.5 mL clean tube. For the triplicate environmental samples collected to serve as “processing positive controls”, 10 µL of the pooled RNA eluate from the PED+ farm samples was spiked into the supernatant. This was completed to ensure that both the downstream RNA isolation processing and environmental materials associated with the samples did not impact PEDV RNA detection. The sample supernatants were frozen at -80°C until RNA isolation could be completed in batches of 96.

The *Quick*-DNA/RNA Viral 96 Kit (Zymo Research Corporation) was used to isolate RNA according to the manufacturer’s instructions, with the following modifications: 1) Viral DNA/RNA Buffer was left on the sample for 10 minutes to allow for an adequate lysis period for environmental samples, 2) centrifuge steps were done at 3,250 rpm for 10 minutes, 3) a five-minute dry spin at 3,250 rpm was done before RNA elution, and 4) elution volume was increased to 18 µL. DNA digestion was performed during the isolation as recommended by the manufacturer to ensure pure RNA. Eluates were kept in 96-well plate format, sealed with a foil seal, and frozen at -80°C until further use.

To serve as a “RT-qPCR positive control,” fresh piglet intestines from PEDV+ diagnostic pathology cases were clipped to make six 50 mg sections of ileum. RNA was isolated from each ileum clipping using the *Quick*-RNA MiniPrep Plus Kit (Zymo Research Corporation) according to the manufacturer’s directions with the following modifications: 1) tissue clippings were each submerged in 600 µL of PBS without calcium and magnesium and homogenized using 2.0 mm lysis beads in a high-speed bead-beater for three rounds of 20-second homogenization and one minute cooling on ice, 2) 60 µL PK buffer and 30 µL Proteinase K were added to each homogenized sample, allowed to incubate at room temperature for 30 minutes, and were centrifuged at 8,000 rpm five minutes to pellet any remaining tissue, and 3) RNA was eluted in two separate 50 µL elutions. The first and second eluates from all six samples were pooled respectively, divided into 20 µL aliquots, and stored at -80°C until further use.

#### 2.3.2 RT-qPCR

An 87 bp fragment of the PEDV N-gene was amplified and measured by RT-qPCR using the Go-Taq Enviro RT-qPCR System (Promega Madison, WI, USA). Each 20 µL reaction consisted of 10 µL of 2X Go-Taq Enviro Master Mix, 0.4 µL Go-Script Enzyme Mix, 0.8 µL of 10 µM forward primer (5’ - GAATTCCCAAGGGCGAAAAT - 3’) (Ortiz et al., 2023), 0.8 µL of 10 µM reverse primer (5’ - TTTTCGACAAATTCCGCATCT - 3’) (Ortiz et al., 2023), 0.2 µL of 10 µM probe (5’ - FAM-CGTAGCAGCTTGCTTCGGACCCA-BHQ1 - 3’) (Ortiz et al., 2023), 5.8 µL of nuclease-free water, and 2 µL of isolated RNA. Additionally, every 96-well-plate of RT-qPCR included two controls: 1) a non-template control consisting of 2 µL of DNA/RNA free water, and 2) a “RT-qPCR positive control” consisting of 2 µL of RNA isolated from the intestines of a PEDV+ piglet. All RT-qPCR reactions were run on the QuantStudio 5 (Applied Biosystems, Waltham, MA, USA) with the following cycling conditions: 45°C for 15 minutes, 95°C for 2 minutes, and 40 cycles of 95°C for 15 seconds and 60°C for 1 minute. RT-qPCR results are reported as quantification cycle (Cq) values. When interpreting the resulting Cq value, an inverse relationship applies: a lower Cq value indicates a higher starting quantity of PEDV RNA in the sample, whereas a higher Cq indicates a lower starting quantity of PEDV RNA.

To confirm the accuracy of the in-house PEDV RT-qPCR assay, 10% of samples (n=254) across different truck types, collection periods, and collection sites (e.g. back-of-trailer, tire, and cabin) were additionally tested for PEDV, TGEV, and PDCoV using a commercial triplex kit (Indical Bioscience, Leipzig, Germany). Each 25 µL reaction consisted of 18 µL virotype® Mix +IC(TAMRA)-RNA, 2 µL virotype® PEDV/TGEV/PDCoV Primers/Probes, and 5 µL isolated RNA and were run on the QuantStudio 5 (Applied Biosystems) under the following cycling conditions: 50°C for 10 minutes, 95°C for 2 minutes, 40 cycles of 95°C for 5 seconds and 60°C for 30 seconds. Every 96-well-plate of RT-qPCR with the commercial triplex included two controls: 1) a non-template control consisting of 5 µL of DNA/RNA free water, and 2) a positive control consisting of 5 µL of the Indical virotype PEDV/TGEV/PDCoV positive control.

### 2.4 Statistical Analysis

While the commercial (Indical) assay has clear interpretation guidelines (Cq ≥ 35 should be considered negative; Cq < 35 should be considered positive), we did not assume those guidelines would be appropriate for the results generated with the in-house assay. Thus, to determine an appropriate cutoff value for determining PEDV RNA positive and negative samples with the in-house assay, two separate approaches were carried out using the Cq values from the samples processed with both the in-house and commercial assay (n=254). Firstly, using data from the custom assay, the receiver operating characteristic (ROC) curve was plotted and the resulting area under the curve (AUC) values were determined for each potential cutoff value (every integer from 25 to 36) using the commercial assay results with a cutoff of 35 as the reference. Secondly, the data was assumed to fit a Weibull distribution using a mixed models procedure. A No-U-turn sampler (NUTS) was used to estimate the posterior distribution of Cq values for positive samples to obtain the 95% credible interval. An agreement between these two methods was used to determine the cutoff value.

Using the statistically determined Cq threshold of 32 established for the in-house assay, study samples (n=2,004) were either deemed positive (≤32) or negative (>32). Statistical analyses were completed using both positive/negative status and Cq value to assess the impact of study variables including truck type, disinfection status (e.g. before or after disinfection), disinfectant used, swab collection site, and month collected on the detection of PEDV RNA. For sample collection duplicates (n=190), Cq values were averaged. Any sample with a Cq>40 was omitted from the analysis (Supplementary Table 2). Summary statistics were calculated for the positive/negative status of samples. For Cq value comparisons, the Shapiro-Wilk test was used to determine normality. For normal data sets, pooled t-tests and Tukey–Kramer HSD tests were used with an ∝ = 0.05. For non-normal data sets, the Wilcoxon signed-rank test and Dunn tests with a Bonferroni correction were performed with an ∝ = 0.05 to determine data significance. Details on the tests completed for each comparison are given in Supplemental Table 1.

## 3. Results

### 3.1 Cq cutoff

The Cq cutoff value for determining positive and negative samples needed to be appropriate for the sensitivity level of the in-house RT-qPCR assay developed for this study. The maximum AUC of 0.8077 was found when using the cutoff Cq value of 33. The second-best AUC of 0.7974 was observed using a cutoff Cq value of 32. Both AUC values are considered “acceptable” (Nahm et al., 2022). The 95% credible interval for Cq values of positive samples determined using the NUTS method was 31.5-32.8. Based on the combined results from both methods, a Cq of ≤32 was selected as the optimal cutoff for classifying a sample as PEDV RNA positive using the custom RT-qPCR assay (Table 2).

**Table 2.** Percentages of samples positive for PEDV RNA based on the in-house RT-qPCR results and a Cq cutoff value of ≤32.

| TRUCK<br>TYPE | DISINFECTION STATUS |  |  |  |  |  |  |
| --- | --- | --- | --- | --- | --- | --- | --- |
|  | *Before Disinfection* |  |  | *After Disinfection* |  |  |  |
|  | *Location of sampling* |  |  | *Advanced Hydrogen Peroxide* |  | *Quaternary Ammonium/Glutaraldehyde* |  |
|  | **Cabin** | **Tire** | **Back-of-Trailer** | **1:40** | **1:32** | **1:128** | **1:100** |
| **PED+** | 50.56% (45/89) | NA | 48.86% (43/88) | NA | NA | 25.00% (7/28) | 36.96% (17/46) |
| **PED-** | 47.25% (43/91) | NA | 41.57% (37/89) | 41.67% (5/12) | 43.75% (7/16) | 23.81% (5/21) | 16.67% (3/18) |
| **Feed** | 16.32% (31/190) | 7.60% (13/171) | NA | NA | NA | 26.92% (23/78) | 3.19% (3/94) |
| **Market** | 87.29% (103/118) | NA | 79.17% (95/120) | 100% (26/26) | 97.22% (35/36) | 70.37% (19/27) | 84.84% (28/33) |
| **Crew** | 67.65% (23/34) | 30.00% (9/30) | NA | NA | NA | NA | NA |

\*NA- Data not available as such samples were not collected from that vehicle type
\*Values are reported as follows: Percentage of PEDV RNA positive samples (number of positive samples/total number of samples).
\*Feed truck samples collected after disinfection were only collected from the tire. PED+, PED-, and market truck samples after disinfection were only collected from the back of the trailer.

### 3.2 Prevalence of PEDV RNA on trucks

A total of 387 individual cabins, trailers, and tires (587 total cabins, trailers, and tires sampled due to repeat visits) were included in the study. The aim was to sample 30% of the production company fleet’s trucks for each truck type; based on availability, more than the 30% of each truck type was sampled for all except feed trucks (27%). The number of unique trucks included in the study per sample type is as follows: crew, 14 cabins/tires; feed, 52 trailers and 63 cabins; pigs-to-market, 72 trailers and 48 cabins; pigs-to-farm, 80 trailers and 44 cabins (PED+ and PEV-combined). From these trucks, 2,004 environmental samples were collected and processed for PEDV RNA detection using RT-qPCR. A total of 92.07% of the “processing positive control” samples were deemed PEDV RNA positive, verifying that the sampling approach, storage, and RNA isolation procedures were appropriate for isolating PEDV RNA. Additionally, 190 samples were collection sample duplicates; Cq values were averaged before further analysis. Of the remaining 1,650 samples, 1,455 samples had detectable PEDV RNA (Cq < 40, 88.18%) and 625 were deemed positive for PEDV RNA (Cq ≤32, 37.88%).

Market trucks had the lowest median Cq value as well as the shortest median wash time of 15 minutes (*pers. obs.*) and the highest percentage of positive results for samples collected before disinfection (Figure 2, Table 2, and Table 3). The highest Cq values and lowest percentage of positive results were observed from feed trucks (Figure 2, Table 2, and Table 3). PED+ and PED-pigs-to-farm truck median Cq values were within 1.40 cycles of each other when comparing samples collected before disinfection, as well as comparing samples disinfected with QAG (Table 3). PED+ pigs-to-farm trucks had a median wash time of 60 minutes, while 45 minutes was the median wash time for PED-pigs-to-farm trucks (*pers. obs*.).

**Figure 2.**
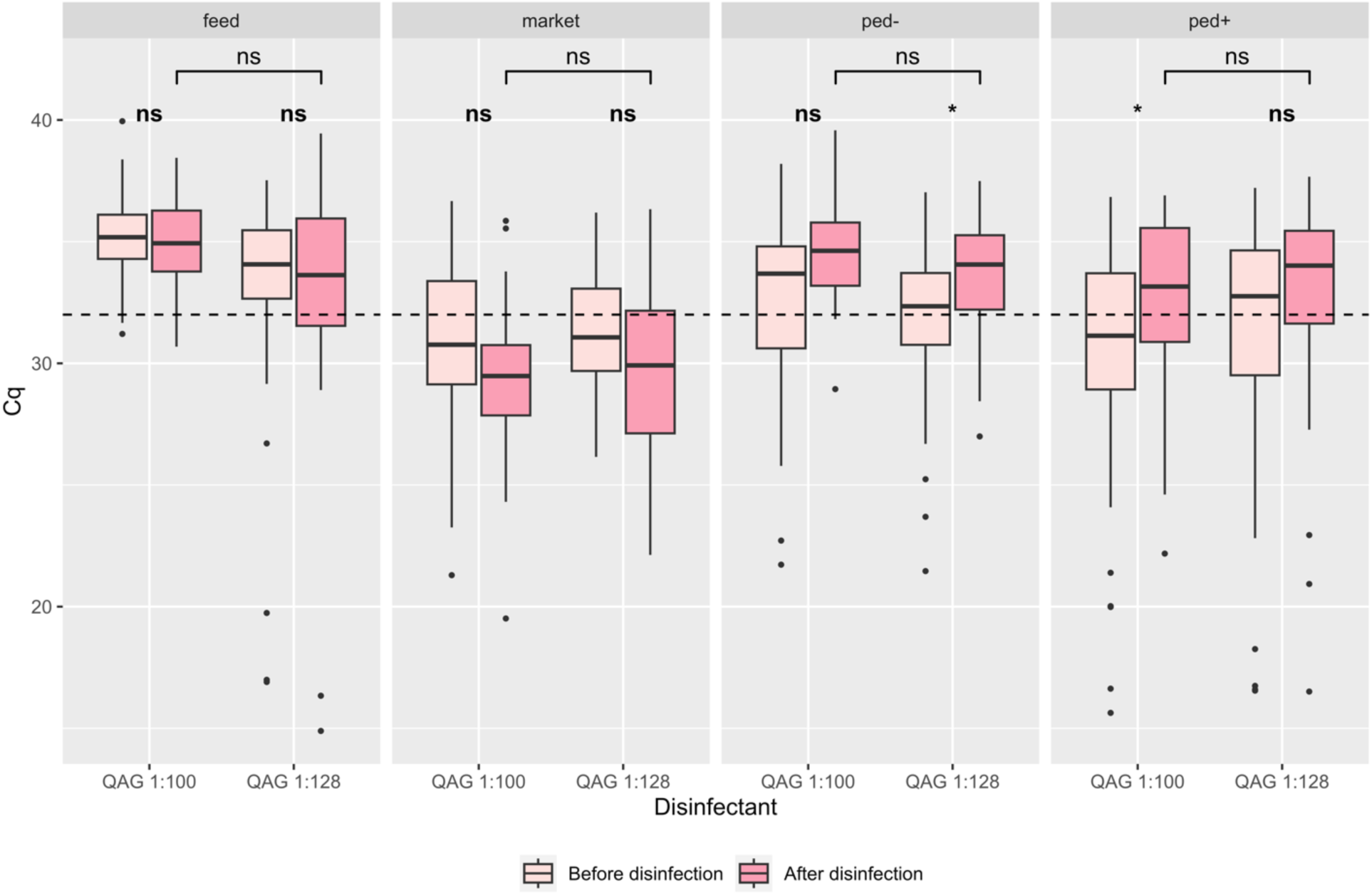
Comparing disinfectant efficacy in removing PEDV RNA from vehicle exteriors (back-of-trailer and tire samples) across truck types (separated by facet). Light pink box and whisker plots show the Cq values for samples before disinfection. Darker pink box and whisker plots show the Cq values for samples after disinfection. Lines indicate the median; the box indicates the 1^st^ and 3^rd^ quartiles; and the whiskers indicate the 1.5 x interquartile range. Individual dots represent outliers. The dashed horizontal line spanning all plots denotes a Cq value of 32, used as the threshold for determining PEDV positive (<32) and negative (≥32) samples. Abbreviations are as follows: Cq, quantification cycle; QAG, Quaternary Ammonium, and Glutaraldehyde; ns, no significance. *p<0.05 (Supplementary Table 1).

**Table 3.**
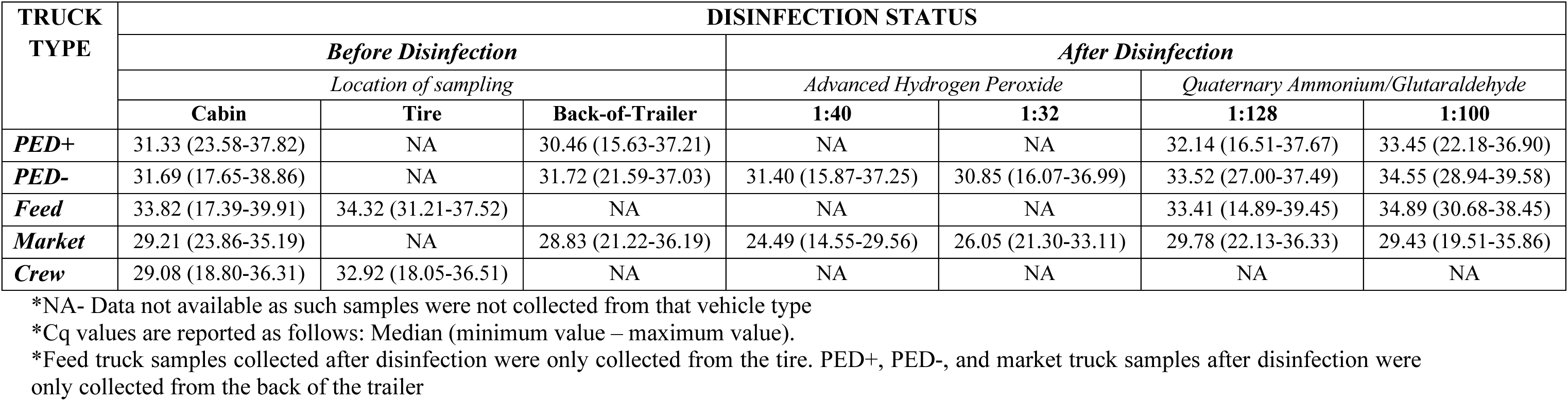
Summary statistics for Cq values obtained from the in-house RT-qPCR results based on truck type, disinfection status, sampling location, and disinfectant.

### 3.3 Effect of disinfectants on PEDV RNA concentration

When considering all back-of-trailer and tire samples and using Cq value as an assessment of cleanliness (i.e., more or less detectable PEDV RNA), neither disinfectant at the high nor low concentration left the trailers/tires significantly cleaner after disinfection (Figure 3, Supplementary Table 1). This is also observed when considering back-of-trailer and tire samples separately (Supplementary Figure 1 and Supplementary Figure 2, respectively). PED-pigs-to-farm trailers and pigs-to-market trailers were subjected to all four disinfection options; only QAG 1:128 left the PED-pigs-to-farm trailers significantly cleaner after disinfection (Figure 4A, p=0.024) while neither disinfectant yielded significantly cleaner market trailers (Figure 4B, p>0.05). Moreover, AHP 1:32 left market truck trailers significantly more contaminated with PEDV RNA after disinfection (Figure 4B, p=0.0197). When considering after disinfection samples from each truck type, the Cq values observed from higher and lower disinfectant concentrations were never significantly different from each other (Figure 2, Figure 4, Supplementary Table 1). Additionally, when looking at market trailers after disinfection, the Cq values observed from trailers disinfected with AHP were significantly lower than what was observed for trailers disinfected with QAG (Supplementary Table 1).

**Figure 3.**
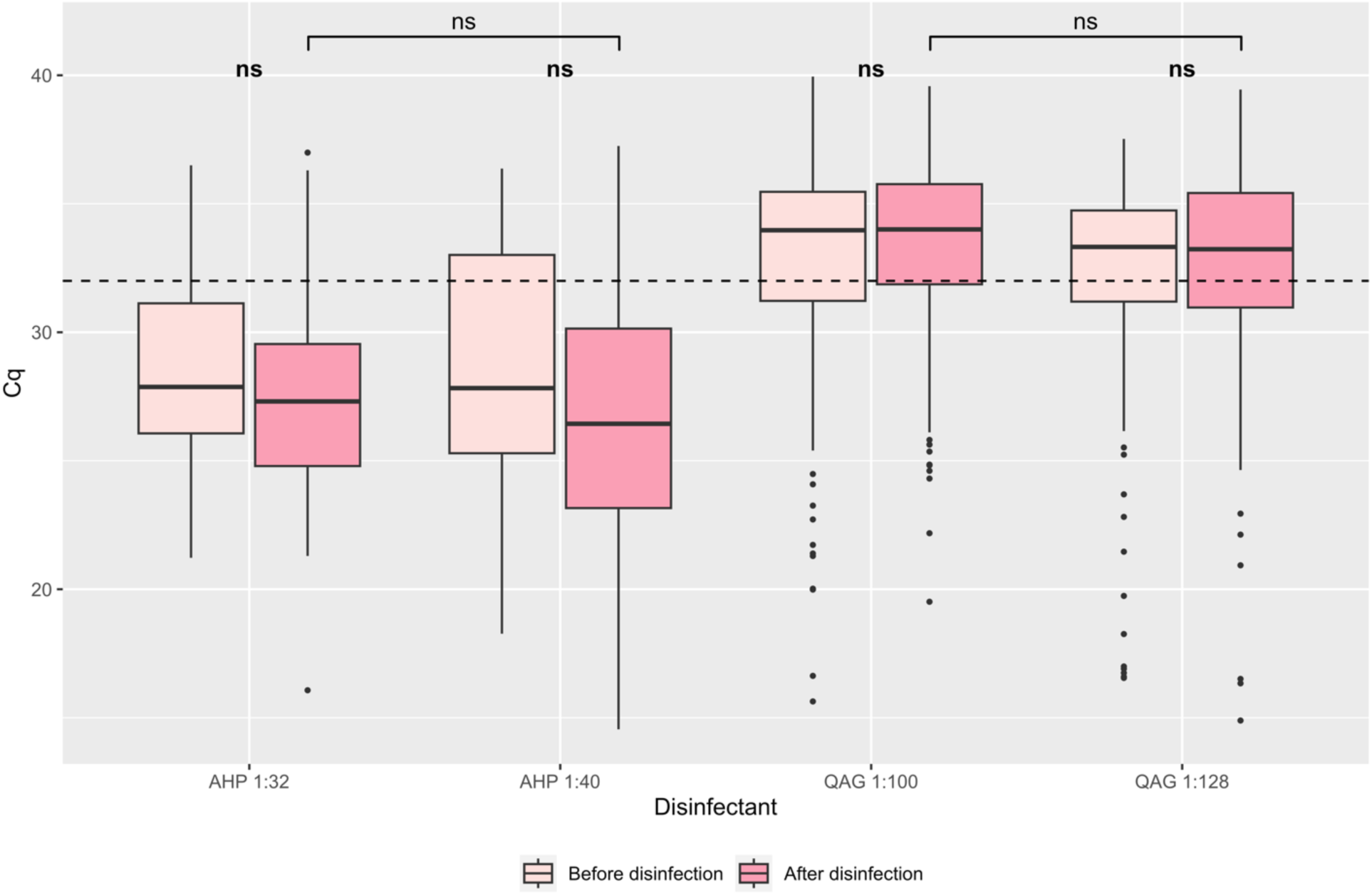
Comparing disinfectant efficacy in removing PEDV RNA from vehicle exteriors (back-of-trailer and tire samples) across pigs-to-farm, pigs-to-market, and feed trucks. Light pink box and whisker plots show the Cq values for combined back-of-trailer and tire samples before disinfection. Darker pink box and whisker plots show the Cq values for combined back-of-trailer and tire samples after disinfection. Lines indicate the median; the box indicates the 1^st^ and 3^rd^ quartiles; and the whiskers indicate the 1.5 x interquartile range. Individual dots represent outliers. The dashed horizontal line spanning all plots denotes a Cq value of 32, used as the threshold for determining PEDV positive (<32) and negative (≥32) samples. Abbreviations are as follows: Cq, quantification cycle; AHP, Accelerated Hydrogen Peroxide; QAG, Quaternary Ammonium, and Glutaraldehyde; ns, no significance (Supplementary Table 1).

**Figure 4.**
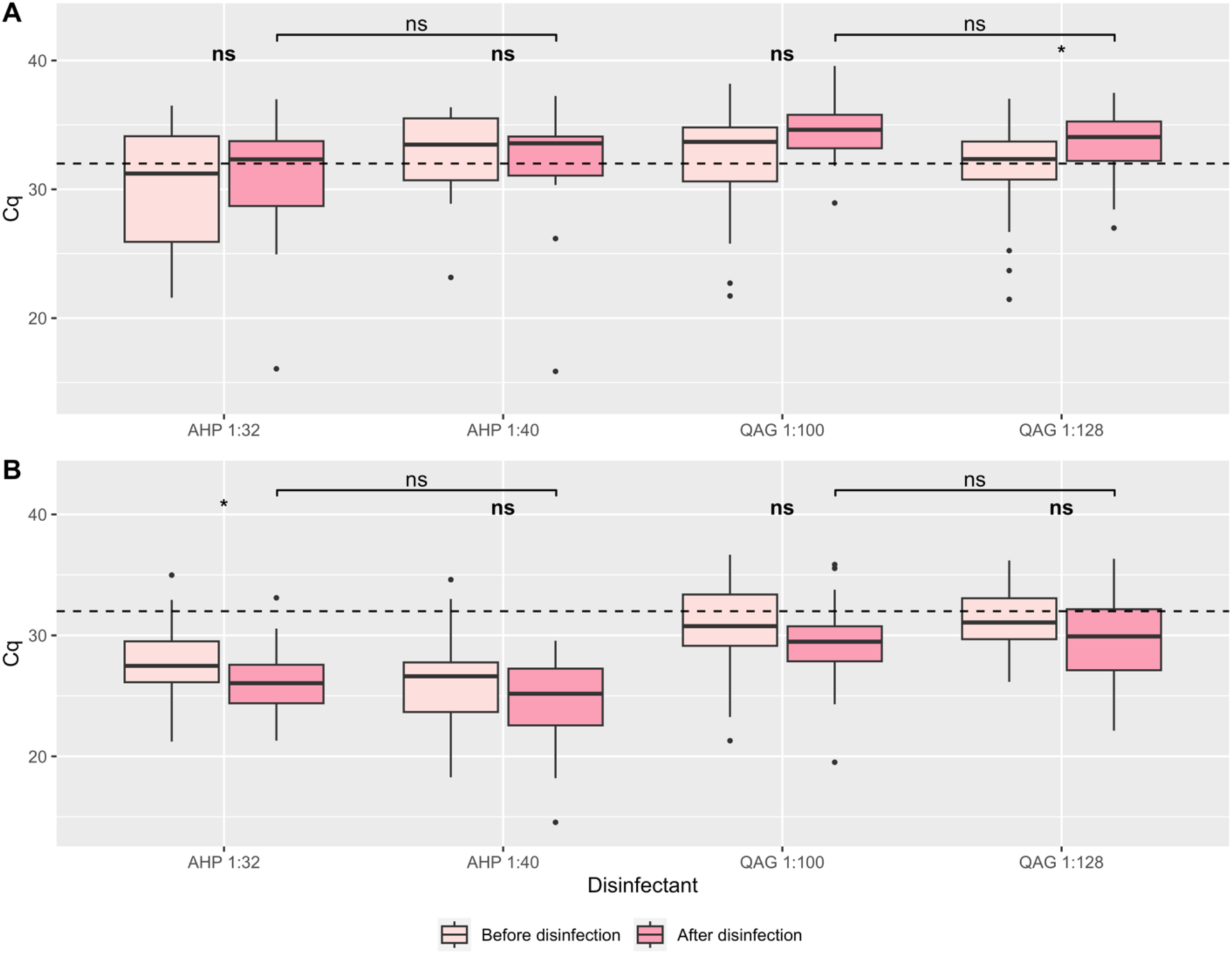
Comparing disinfectant efficacy across (A) back-of-trailer samples from PED-pigs-to-farm trucks and (B) back-of-trailer samples from pigs-to-market trucks. Light pink box and whisker plots show the Cq values for samples before disinfection. Darker pink box and whisker plots show the Cq values for samples after disinfection. Lines indicate the median; the box indicates the 1^st^ and 3^rd^ quartiles; and the whiskers indicate the 1.5 x interquartile range. Individual dots represent outliers. The dashed horizontal lines spanning all plots denote a Cq value of 32, used as the threshold for determining PEDV positive (<32) and negative (≥32) samples. Abbreviations are as follows: Cq, quantification cycle; AHP, Accelerated Hydrogen Peroxide; QAG, Quaternary Ammonium, and Glutaraldehyde; ns, no significance. *p<0.05 (Supplementary Table 1).

Test strips were used to determine the concentration of residual disinfectants after they were applied PED- and market trucks. Of the 49 trucks disinfected with AHP 1:32 and evaluated with test strips, seven had a reading of 1:32 or higher. Of the 33 trucks disinfected with AHP 1:40 and tested, four had a reading of 1:32 or higher. Of the 28 trucks disinfected with QAG 1:100 and tested, five had a reading of 1:100. Lastly, of the 44 trucks disinfected with QAG 1:128, 12 had a reading of 0.5 or higher (equivalent to 1:200). In the majority of cases, the disinfectant remaining on the truck was lower than the optimal concentration for effective disinfection. Notably, all of our test samples were collected from the back of the trailer.

### 3.4 Impact of collection site on PEDV RNA detection

Three different collection sites were sampled for this study: 1) back-of-trailer for the live-animal vehicles, 2) tires for the crew and feed trucks, and 3) the truck cabins for all vehicle types (Figure 1). Market trucks were far more contaminated with PEDV RNA with the lowest median Cq values for the cabin and the back-of-trailer samples while crew truck tires and cabins had lower Cq values compared to feed trucks (Table 3, Figure 5A). When considering all pre-disinfection data, each of the three collection sites were significantly different from each other; the tire samples had a significantly higher Cq values than the cabin samples and the back-of-trailer samples had a significantly lower Cq values than the cabins (Figure 5B, p<0.0001). For each truck type, the cabins had a higher percentage of samples deemed PEDV RNA positive compared to the corresponding back-of-trailer or tire samples (Table 2). Additionally, for all live-haul vehicles, there was not a significant difference between the Cq values of the cabin and the back-of-trailer samples (Figure 5A, Supplementary Table 1).

**Figure 5.**
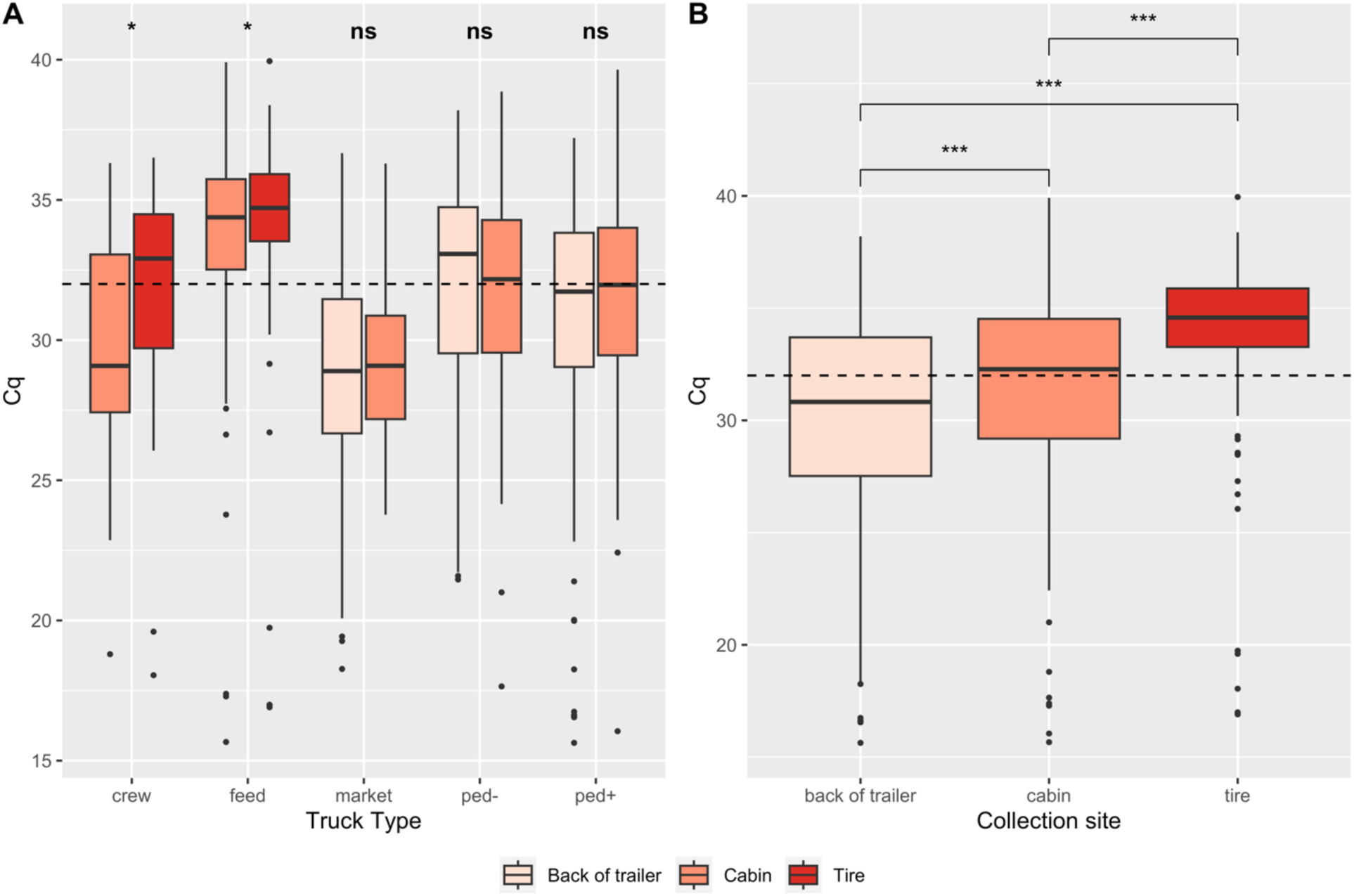
Comparing pre-disinfection Cq values across (A) collection sites with respect to truck type and (B) collection sites across all truck types. Light peach box and whisker plots show the Cq values for samples collected from the back-of-trailer. Orange box and whisker plots show the Cq values for samples collected from the cabin. Red box and whisker plots show the Cq for samples collected from the tire. Lines indicate the median; the box indicates the 1^st^ and 3^rd^ quartiles; and the whiskers indicate the 1.5 x interquartile range. Individual dots represent outliers. The dashed horizontal line spanning both plots denotes a Cq value of 32, used as the threshold for determining PEDV positive (<32) and negative (≥32) samples. Abbreviations are as follows: Cq, quantification cycle; ns, no significance. Wilcoxon tests used to determine statistical significance; *p<0.05, ***p≤0.0001

### 3.5 Impact of seasonality

Samples were collected during 14 bi-weekly collection periods between November 2023 and May 2024. Collection weeks one through seven took place during the winter period (November 2023-February-2024) and collection weeks eight through 14 were during the spring period (March 2024-May-2024). Four (57%) of the winter collection weeks had higher median Cq values after disinfection, whereas only three (42%) of the spring collection weeks had this result. However, the median Cq values before disinfection for the winter collections were typically lower than the spring collections (Figure 6). Although Cq values for back-of-trailer and tire samples collected after disinfection were not significantly different between seasons, the Cq values observed for back-of-trailer and tire samples collected before disinfection were significantly lower in winter than in spring (p=0.016).

**Figure 6.**
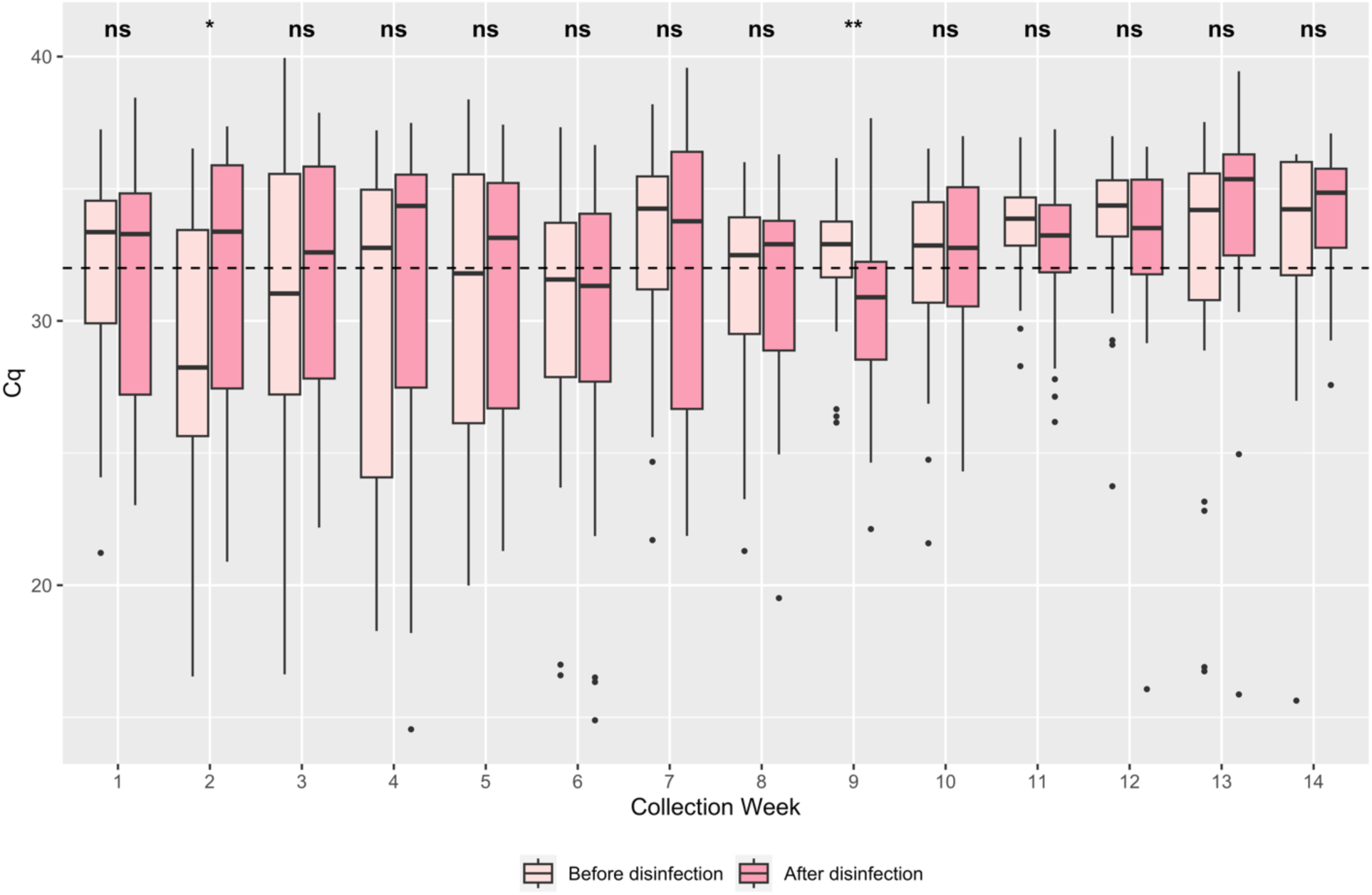
Comparing disinfectant efficacy in removing PEDV RNA from pigs-to-farm, pigs-to-market, and feed truck exteriors (back-of-trailer and tire samples) across collection weeks. Seasonality is as follows: Winter, November-February, weeks 1-7; Spring, March-May, weeks 8-14. Light pink box and whisker plots show the Cq values for samples before disinfection. Darker pink box and whisker plots show the Cq values for samples after disinfection. Lines indicate the median; the box indicates the 1^st^ and 3^rd^ quartiles; and the whiskers indicate the 1.5 x interquartile range. Individual dots represent outliers. The dashed horizontal line denotes a Cq value of 32, used as the threshold for determining PEDV positive (<32) and negative (≥32) samples. Abbreviations are as follows: Cq, quantification cycle; ns, no significance. Wilcoxon tests used to determine statistical significance; *p<0.05, **p<0.001.

## 4. Discussion

By collecting samples from swine industry vehicles and trailers and utilizing an in-house RT-qPCR assay to detect PEDV RNA, we were able to examine the contamination level of vehicles with PEDV RNA before and after disinfection across different vehicle types when cleaned with different types and concentrations of disinfectants. The in-house assay, which is a more cost-effective alternative to commercial assays, was specifically optimized for use with environmental vehicle samples, which contain less genetic material compared to samples taken directly from organisms having clinical viral symptoms. We demonstrated that market trucks were a) the most contaminated with PEDV RNA, b) the most likely truck type to still be physically dirty after washing (*pers. obs.*), and c) washed for the shortest amount of time (*pers. obs.*) (Figure 2). This is most likely because these vehicles are only carrying pigs who are exiting the swine production system (e.g. the slaughterhouse) and no longer bear the risk of carrying PEDV to sow farms (e.g. Mannion et al., 2008; Galvis and Machado, 2024). Pigs-to-farm trailers, which carry pigs from all production phases before being sent to the market, were more effectively washed and were significantly less likely to be contaminated with PEDV RNA compared to market trucks (Table 3, Figure 2; PED+ pigs-to-farm, p=0.0006, PED-pigs-to-farm, p<0.0001) (Galvis and Machado, 2024). Pigs-to-farm trucks from PED+ and PED-farms were never significantly different from each other, despite being deliberately separated (e.g. washed at different C&D stations, driven by separate personnel, and sent to only farms with the same PEDV status) (Table 3, Supplementary Table 1). RNA can persist in environments which facilitate the stability of nucleic acids, such as in the presence of organic material, so it is plausible that trucks leaving PED-farms are testing positive due to residual PEDV RNA from previous trips. The most notable discrepancy between the PED+ and PED-pigs-to-farm trucks was that for PED+ farm trucks, 36.96% of the back-of-trailer samples disinfected with QAG 1:100 were PEDV RNA positive while only 16.67% of the back-of-trailer samples disinfected with QAG 1:100 from PED-farms were positive (Table 3, p=0.196). However, when comparing these samples disinfected with QAG 1:128, the percentages of PEDV RNA positive samples become much more comparable (Table 2, p=1.000).

Given there was not a significant difference between the cabin and back-of-trailer pre-disinfection samples for all three truck types, it is likely that contaminants (possibly feces) are moving through fomites (e.g. boots, equipment) between the interior and exterior of the vehicles (Figure 5A, Supplementary Table 1). Since the crew and feed truck cabin samples were significantly more contaminated with PEDV RNA than the respective tire samples in this study, cabins could likely benefit from regular C&D similar to the exterior of the vehicles.

Based on PED-pigs-to-farm and market trucks, which were the only truck types disinfected with both disinfectants, QAG was the more effective disinfectant compared to AHP (Figure 4). QAG is on the approved list of disinfectants for decontaminating PEDV and has been shown to disrupt infectious PEDV in treated samples, but not PEDV RNA (EQSP report 2015; Bowman et al., 2015). QAG was able to lower the percentage of PEDV RNA positive samples from before disinfection to after disinfection in 75% of the scenarios it was used, while AHP was not effective in this manner for either of the truck types it was used with (Table 3). Disinfectants were prepared by production company staff at the beginning of every day and subsequently tested prior to use; our team used test-strips to evaluate the disinfectant concentration after vehicle application. Notably, the test-strip concentrations varied greatly and were typically low compared to the desired concentration. Resulting Cq values were rarely significantly different from each other; only AHP 1:32 saw significant differences in Cq values when comparing across concentration readings (readings of 0 and 1:16, p=0.0083; all other pairs non-significant). The unreliable and low readings may be the consequence of applying disinfectant directly after washing. In such a scenario, the leftover water from washing is still running off the trailer and pooling on the surfaces during the disinfectant application, thus diluting the disinfectant to undetectable levels. The ideal solution for this would be to allow the truck to dry before applying the disinfectant, then allowing it to make contact for the manufacturer’s recommended time; however, these may not be feasible due to company time constraints and the trailer’s vertical surfaces (which the disinfectant cannot soak on *[pers. comm.])*. A possible alternative could be to apply the disinfectant at a high enough concentration that once it mixes with the residual water, it will dilute to a level that will effectively kill viral pathogens (EQSP report 2015). However, this needs to be examined more thoroughly as previous research rarely considers the concentration of the disinfectant after it is applied to a washed vehicle (Bowman et al., 2015; Boniotti et al., 2018).

Even though our feed trucks were relatively free of RNA contamination compared to the other truck types, research has shown that the only surfaces in a feed mill that had detectable RNA from porcine enteric viruses (including PEDV) were from the feed delivery system (Elijah et al., 2021). Previous studies have also shown that similar to our findings, feed delivery truck cabins were likely to be contaminated (Elijah et al., 2021; Houston et al., 2024; Greiner, 2016). Although we could not evaluate whether the drivers for any of the trucks were exiting their vehicles at their destinations (e.g. the slaughterhouse, farm, feed mill), it has been shown that when drivers leave their cabin (or when other personnel step onto the trailers) the risk of contamination increases (Lowe et al., 2014; Greiner, 2016; Houston et al., 2024; Elijah et al., 2021). This could explain why crew truck cabins were significantly more contaminated with PEDV RNA than the tires from the same vehicle, as it is very likely crew members are entering a contaminated area at their destination and then tracking PEDV RNA into their cabin when leaving.

Despite this study’s attempt to maintain consistency by selecting C&D stations operated by the same production company, truck C&D was performed by company personnel and therefore could not be validated across sites; wash times, trailer cleanliness after disinfection, amount of disinfectant applied, and thoroughness of disinfectant application all varied between stations and between station personnel. Additionally, the recommended contact time for disinfectants could not always be followed, especially for vertical surfaces. Companies which manufacture QAG recommend a contact time of 5-12 minutes; however, due to the nature of the C&D stations, post-disinfection samples were often taken less than five minutes after the disinfectant was applied. Another limitation, concerning RT-qPCR results, was that not all of the “processing positive control” samples resulted in Cq values ≤ 32 (which would be classified as PEDV RNA positive). While we would ideally want this to be 100%, these “processing positive controls” still confirmed that the storage solution (DNA/RNA shield; Zymo Research), the RNA isolation kit, and the in-house RT-qPCR were working effectively. Nonetheless, the main limitation of this study is that detecting PEDV RNA using RT-qPCR does not necessarily mean that there is an infectious virus present. Commercial disinfectants are designed to target viral capsids which often leave the RNA intact (Bowman et al., 2015). To study infectivity, the gold-standard approaches include growing the virus in cell culture or an animal model (bioassay) (Puente et al., 2020). Recent research suggests that viability RT-qPCR assays can determine if there is viable virus in a sample (Puente et al., 2020; Balestreri et al., 2024). However, these methods are unfeasible for pork producers due to time, validity, and material restrictions, especially during large outbreaks (Bowman et al., 2015). RT-qPCR allows the industry to determine if PEDV should be an immediate concern for their herds and to quickly address the outbreak.

## 5. Conclusion

The aim of this study was to determine how frequently industry vehicles are contaminated with PEDV RNA before and after C&D. We determined that pigs-to-market trucks are the most likely to be contaminated overall and that pigs-to-farm trucks from PED-farms have similar PEDV RNA loads to pigs-to-farm trucks from PED+ farms. We also identified that the cabins of all five truck types are not significantly cleaner than the exterior of the trucks. Based on these results, vehicle biosecurity measures involving the disinfection of truck interiors, as well as market truck disinfection as a whole, should be evaluated to prevent the spread of PEDV as trucks go back to farms after washing and as personnel exit and enter their cabins during vehicle visits. Additionally, this study identified QAG to be a more effective disinfectant than advanced hydrogen peroxide for decreasing PEDV RNA contamination of swine industry vehicles.

## Ethical statement

The authors confirm the journal’s ethical policies, as noted on the journal’s author guidelines page. Ethics permits were unnecessary since this work did not involve animal sampling or questionnaire data collection by the researchers.

## CRediT authorship contribution statement

TBP: Data curation, formal analysis, investigation, methodology, project administration, validation, visualization, writing – original draft.

KAM: Conceptualization, data curation, funding acquisition, methodology, project administration, resources, supervision, writing – original draft.

GM: Conceptualization, data curation, funding acquisition, methodology, supervision, visualization, writing – review & editing.

MR: Supervision, writing – review & editing.

BSD: Investigation, visualization.

JBF: Conceptualization, data curation, funding acquisition, investigation, methodology, project administration, resources, supervision, writing – original draft.

## Data availability statement

The data supporting this study’s findings are not publicly available and are protected by confidential agreements; therefore, they are not available.

## Acknowledgements

The authors would like to thank the production company that worked with us for this study, including changing disinfectants as requested and allowing us to sample their trucks. We also thank those who assisted in sample collections (Meredith Moss, Makayla Holloman, Ana Castro, Laya Kannan S. Alves, Rubia Tomacheuski, Don Banks, and Eli Parker) and individuals who assisted with wet-laboratory processing (Gabriella Alexander and Mallory Cerak). Additionally, we thank Yang Qu for statistical consulting and Barb Sherry and Shannon Chiera for assisting with the project set-up and processing of tissues.

